# Parallel shifts in flight-height associated with altitude across incipient *Heliconius* species

**DOI:** 10.1101/2023.04.12.536626

**Authors:** David F Rivas-Sánchez, Carlos H. Gantiva-Q, Carolina Pardo-Diaz, Camilo Salazar, Stephen H Montgomery, Richard M Merrill

**Affiliations:** School of Biological Sciences, University of Bristol, 24 Tyndall Avenue, Bristol, UK; Division of Evolutionary Biology, Ludwig-Maximilians-Universität München, Grosshaderner Strasse 2, 82152 Planegg-Martinsried, Germany; Department of Biology, Faculty of Natural Sciences, Universidad del Rosario, Carrera 24 # 63C-69, Bogotá, 111221, Colombia

**Author notes:** **Corresponding authors:** DFRS, SHM, RMM. Contributed equally (Stephen H Montgomery, Richard M Merrill).

**Keywords:** Heliconiini, local adaptation, micro-habitat, parallel evolution

## Abstract

Vertical gradients in microclimate, resource availability and interspecific interactions are thought to underly stratification patterns in tropical insect communities. However, only a few studies have explored the adaptive significance of vertical space use during early population divergence. We analysed flight-height variation across speciation events in *Heliconius* butterflies representing parallel instances of divergence between low and high-altitude populations. We measured flight-height in wild *H. erato venu*s and *H. chestertonii*, lowland and mountain specialists respectively, and found that *H. chestertonii* consistently flies at a lower height. We compared these data with previously published results for *H. e. cyrbia* and *H. himera*, the latter of which flies lower and, like *H. chestertonii*, recently colonised high-altitude, dry forests. We show that these repeated trends largely result from shared patterns of selection across equivalent environments, producing parallel trait-shifts in *H. himera* and *H. chestertonii*. Although our results imply a signature of local adaptation, we did not find an association between resource distribution and flight-height in *H. e. venus* and *H. chestertonii*. We discuss how this pattern may be explained by variation in forest structure and microclimate. Overall, our findings underscore the importance of behavioural adjustments during early divergence mediated by altitude-shifts.

## Introduction

The steep environmental gradients that characterise the upper and lower vegetation layers of tropical forests are a key factor determining species diversity and abundance [1-5]. In arthropods, particular species are consistently found in some vertical strata while being less common in others due to variation in abiotic conditions, behaviour, and the distribution of resources [6, 7]. For example, fruit-feeding and nectar-feeding butterflies, respectively decrease and increase in abundance towards the canopy, ostensibly tracking their food sources [8-10]. This has broad implications for the structure of communities, with consequences for wider trophic interactions [11, 12].

Comparatively, less is known about vertical segregation occurring between closely related populations. Only a handful of studies have examined divergence in the use of vertical space at this level, but these provide a range of evidence implicating flight differences in early divergence. For instance, *Heliconius* flying in separate vertical strata have different wing morphologies [13]. Similarly, the Nymphalid *Archaeoprepona demophon* includes coexisting canopy and undercanopy populations with independent demographic dynamics and low levels of gene-flow [14]. Divergence in resource use, combined with changes in larval behaviour, have also been linked to divergence in *Colobura dirce* [15], which can be split into two separate species based on differences between their larvae: *C. annulata* are gregarious and feed on mature, canopy *Cecropia* leaves while *C. dirce* are solitary and consume undercanopy saplings [15]. These examples suggest a role for vertical space in local adaptation, but the underlying mechanisms remain unclear.

Recently, heritable flight-height differences were demonstrated between the *H. e. cyrbia* and *H. himera* [16], which are distributed along a gradient from wet, low-altitude to dry, high-altitude forest, with a narrow zone of overlap where they hybridise [17]. Flight-height correlates with the distribution of resources in their habitats such that *H. himera* flies lower than *H. e. cyrbia*, tracking the height of hostplants and flowers that are positioned lower at high-altitude [16]. This pattern may result from divergent ecological selection maximising hostplant encounter rates and foraging efficiency across forests with different structures, potentially contributing, alongside other factors [18-22], to the evolution of reproductive isolation.

In the western slope of the Colombian Cordillera Occidental, *H. e. venus* and *H. chestertonii* are distributed along a similar cline (Figure 1 d-e). *H. e. venus* occupies wet forests at sea level, whereas its close relative *H. chestertonii* lives in drier scrub forests in the mountains [23, 24]. Because *H. chestertoniii* is spatially and genetically independent from but ecologically comparable to *H. himera* [25], it can be studied as a convenient replica to explore the relationship between vertical segregation and divergence during the early stages of speciation. Here, we tested whether *H. e. venus* and *H. chestertonii* fly at different heights and if this correlates with the location of larval and adult-stage resources in their respective habitats. Assuming that flight-height patterns constitute adaptive traits, we predicted that flight-height variation between *H. e. venus* and *H. chestertonii* would mirror that between their ecological equivalents *H. e. cyrbia* and *H. himera*.

**Figure 1.**
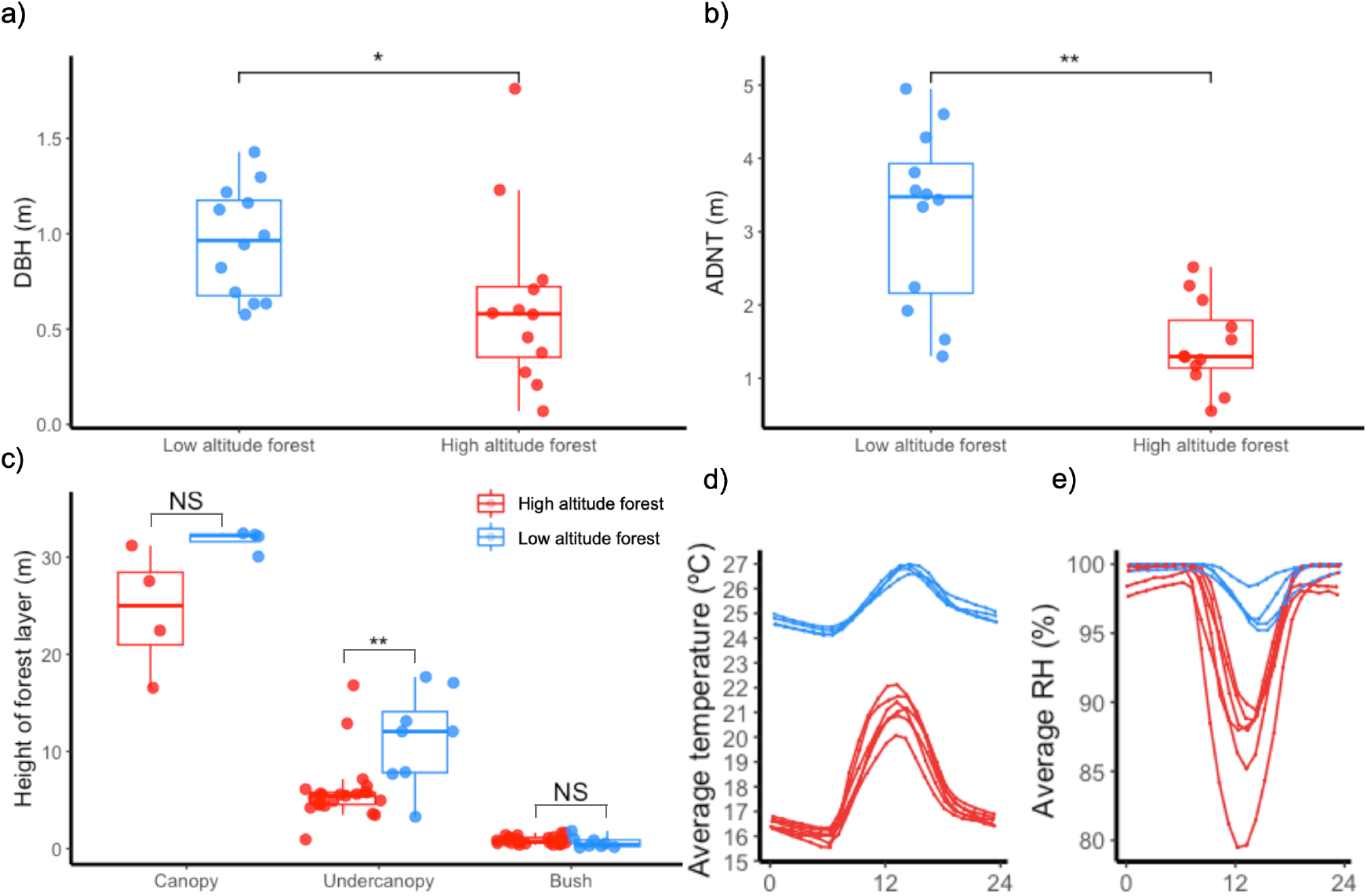
Characteristics of low-altitude (blue) and high-altitude (red) habitats. Comparison of a) diameter at the breast height (DBH), b) average distance between neighbouring trees (ADNT), c) canopy, undercanopy, and shrub heights, d) temperature and e) relative humidity throughout the day. In a-c), box plots show median and upper and lower quartiles, each point is an individual. Significance levels (Wilcoxon-test): NS, not significant. * p<0.05. ** p<0.01. In d-e) (modified from [22]), lines represents microclimate recorded by individual dataloggers deployed in low (blue) or high-altitude (red) forest.

## Methods

### i) Habitat characterisation

We investigated *Heliconius* butterflies in sites representing the habitats of *H. e. venus* (Buenaventura, N03º50’0.04” W77º15’45.1” and La Barra, N03°57.558420 W 77°22.705080) and *H. chestertonii* (Montañitas N03º40’36.3” W76º31’19.4” and El Saladito N03º29’11.5” W76º36’39.8”) in January 2021. At each of these, we characterised the local forest structure by measuring the trunk diameter at breast height (m) and the average distance (m) between neighbouring trees of common tree species. Additionally, we used clinometers and measuring tapes to estimate the height (m) of the canopy, the undercanopy, and the bush.

### ii) Vertical distribution of resources

We explored whether the distribution of adult and larval resources differed between habitats. To do this, we measured the height (m) of all the *Passiflora* (Passifloraceae) hostplants that were visually identified along the trails used to collect butterflies. These plants were subsequently identified to the species level [26, 27] and filtered to include only species known to be used by species in the *erato* complex [28, 29], to which our focal species belong. We also measured the height (m) of the flowering plants that butterflies were observed feeding from at least once. Note that these methods differ slightly from Dell’Aglio, Mena [16], where plants were identified by following butterflies until they landed on a flower or hostplant. Due to the trail and forest structure, this approach was not possible here.

### iii) Patterns of flight-height divergence

We measured the flight-height (m) of *H. e. venus* (n=33) and *H. chestertonii* (n=39) at the point where individuals were first encountered using measuring tapes. Individuals were collected, sexed and marked, ensuring no repeated observations. To quantify levels of parallelism in flight-height between butterflies from independent, ecologically equivalent pairs, we included flight-height measures of *H. e. cyrbia* (n=78) and *H. himera* (n=86) in our dataset, available in Dell’Aglio, Mena [16].

### iv) Statistical analyses

To test for significant differences in the diameter at the breast height, the average distance between neighbouring trees, and the height of different forest layers between habitats, we used Wilcoxon rank sum tests implemented in R [30]. We also modelled variation in the height of hostplants and flowers separately using linear mixed-effect models (implemented in *lme4* [31]) with “habitat” as a fixed factor and “site” as a random effect. This was followed by Wald χ^2^ tests (implemented with the *Anova* function in the package *CAR* [32]). To analyse flight in *H. e. venus* and *H. chestertonii*, we used linear-mixed models where variation in flight-height was explained by the fixed factors “subspecies” and “sex”, as well as interactions and the random factor “site”. We subsequently used the *drop1* function (base R) to remove model terms unless their deletion was detectable via likelihood-tests. This resulted in a reduced model with “subspecies” as the only main factor.

Finally, to test for parallel shifts in flight-height across independent divergence events, we used a variance partitioning approach [33] combining data on *H. e. venus, H. chestertonii, H. e. cyrbia* and *H. himera*. In detail, our model included the main factors “habitat” (similarity of selection regimes across independent divergence events) and “locality” (variation in flight-height that responds to the unique properties of each divergence), as well as the “habitat*locality” interaction (inconsistency in magnitude and/or direction of divergence patterns of the two pairs of species). We assessed the significance of each of these terms via likelihood (*drop* 1 function) and Wald χ^2^ tests between alternative models. We then estimated semi-partial R^2^ coefficients (*r2beta* function, Kenward Roger method in the R package r2glmm [34]) in our final model, providing a rough approximation of the proportion of variance explained by each term.

## Results

The habitats of *H. e. venus* and *H. chestertonii* broadly differed (Figure 1a-c). In particular, the trunk diameter at the breast height (W=32, p<0.05) and the distance between neighbouring trees (W=15, p<0.01) were both significantly higher in the habitat of *H. e. venus*. The height of the canopy and shrub forest layers were not significantly different between the two forest types, but the undercanopy was significantly shorter in the habitat of *H. chestertonii* (W=30, p<0.01). We were unable to confirm significant, between-habitat differences in the vertical distribution of hostplants (n=37, χ_1_^2^=0.3949; p=0.5297) or flower resources (n=107, χ_1_^2^=0.0096; p=0.9219) (Figure 2a-b).

**Figure 2.**
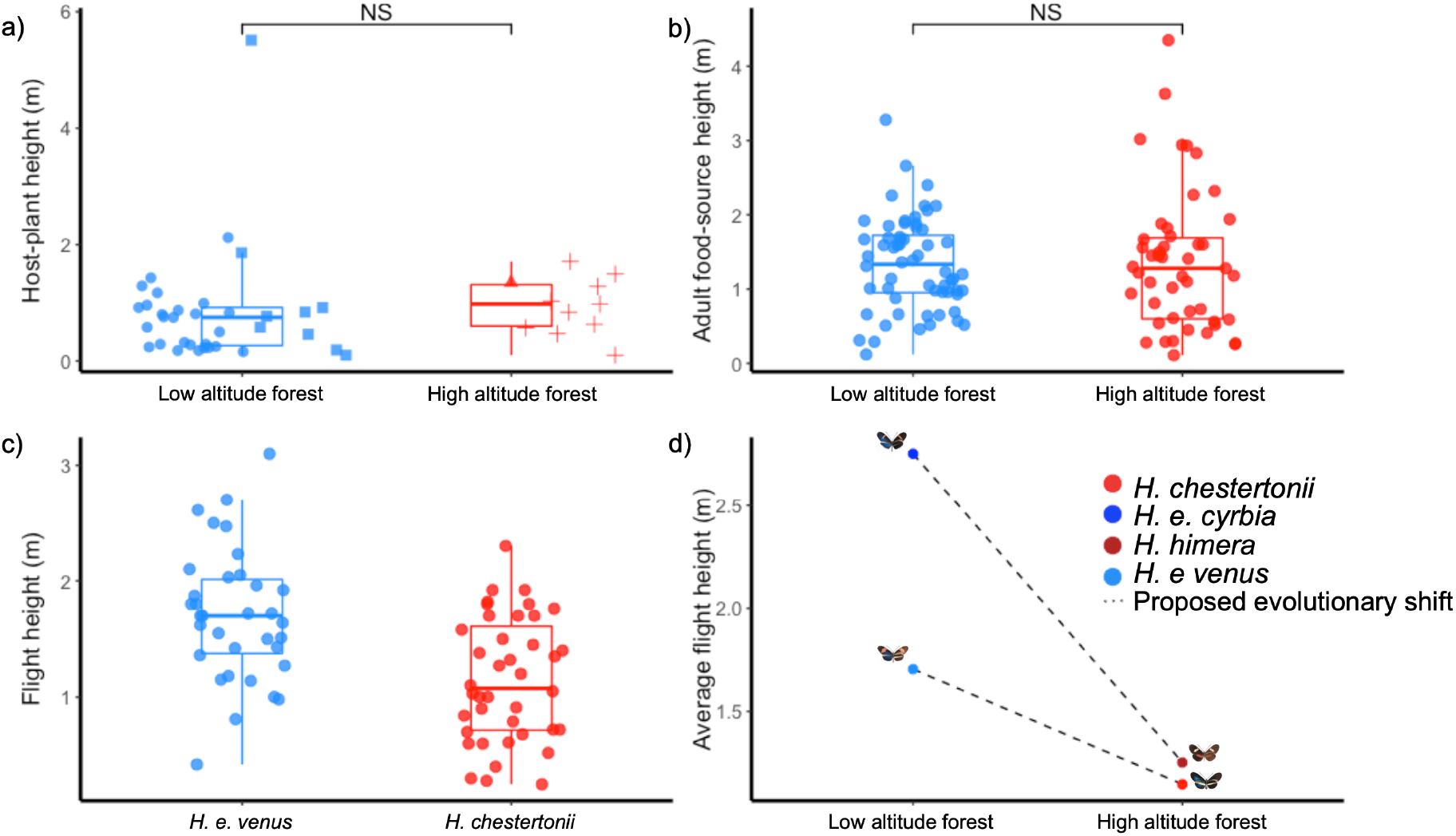
Comparisons in resource distribution and flying-heights in *Heliconius* species. a) Hostplants, b) Adult-food sources, c) Flight-heights and d) Parallelism between different clines. For a) circles are *Passiflora auriculata*, squares are *P. micropetala*, pluses are *P. suberos*a and triangles *P. biflora*. For a-c) *H. e. venus* is blue and *H. chestertonii* is red. For d). *H. e. venus* is blue and *H. chestertonii* is light red, *H. e. cyrbia* is dodger blue and *H. himera* is firebrick. Box plots show median and upper and lower quartiles. Significance levels (Wilcoxon-test): NS, not significant.

On average, *H. chestertonii* flew ∼0.5m lower than *H. e. venus* (n=74, χ_1_^2^=8.8601; p<0.01; Figure 2c). Moreover, likelihood-ratio comparisons between models combining data from *H. e. venus, H. chestertonii, H. e. cyrbia* and *H. himera* showed that the “habitat” (n=240, χ_1_^2^=21.725; p<0.001) and “locality” (n=240, χ_1_^2^=3.894; p<0.05) effects were significant, but not their interaction, and “habitat” explained above 50% of the variance in flight-height (Table 1). This suggests a close correspondence between the patterns of flight-height divergence in *H. e. venus*-*H. chestertonii* and *H. e. cyrbia*-*H. himera*, despite the larger magnitude of difference (∼1.5m) in *H. e. cyrbia*-*H. himera* (Figure 2d).

**Table 1.**
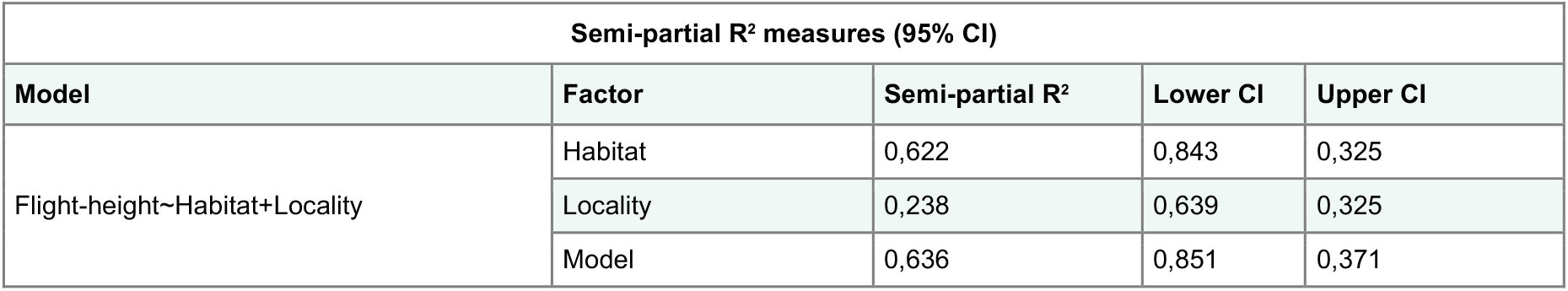
Semi-partial R^2^ measures of terms in model explaining variation in flight-height.

| Semi-partial R<sup>2</sup> measures (95% CI) |  |  |  |  |
| --- | --- | --- | --- | --- |
| Model | Factor | Semi-partial R<sup>2</sup> | Lower CI | Upper CI |
| Flight-height~Habitat+Locality | Habitat | 0,622 | 0,843 | 0,325 |
|  | Locality | 0,238 | 0,639 | 0,325 |
|  | Model | 0,636 | 0,851 | 0,371 |

## Discussion

We leveraged separate events of divergence across ecologically equivalent *Heliconius* pairs in Colombia and Ecuador to explore local adaptation in the context of altitude shifts. We demonstrate that the mountain specialist *H. chestertonii* flies at a lower height compared to the ancestral-proxy, low-altitude *H. e. venus*, mirroring a pattern formerly reported in *H. himera* and *H. e. cyrbia* [16]. This strongly implies a role for related sources of divergent selection driving local adaptation in flight-height across habitat types. However, we were unable to formally link shifts in flight-height in *H. chestertonii* to the distribution of plant resources.

Adaptation can be inferred when parallel phenotypic differentiation occurs across replicate populations in similar environments [33, 35, 36]. The derived populations of *H. chestertonii* and *H. himera* independently colonised similar mountainous habitats [22], as evidenced from genomic data that isolates them in separate clusters [23, 25]. Although we measured flight-height in wild individuals, the differences we found are likely heritable, as implied in common garden experiments with *H. e. cyrbia* and *H. e. himera* [16], and therefore, subject to selection. Indeed, our tests for parallelism suggest these independent shifts in flying behaviour are largely driven by shared sources of divergent selection, rather than stochastic evolutionary processes (e.g., genetic drift) or unique selection regimes [33, 36, 37] and thus potentially indicate repeated local adaptation (e. g. [38-42]).

In contrast to *H. e. cyrbia* and *H. himera* [16], we found no evidence for an association between flight-height in *H. e. venus* and *H. chestertonii* and the vertical distribution of resource plants. In other species, such correlations are interpreted as an indication that butterflies fly at heights that optimise encounter rates with food sources [8, 9, 43, 44]. *P. auriculata* was the only suitable hostplant in *H. e. venus* habitat, whereas we found *P. biflora* and *P. suberosa* in *H. chestertonii* habitat.

These three *Passiflora* belong to the *Decaloba* subgenus and grow to modest dimensions [26], likely explaining their similar heights. This lack of differentiation conflicts with the hypothesis that flight-height is selected on to maximise foraging and oviposition opportunities. Interestingly, in the low-altitude, *H. e. venus* site La Barra, we found plants of a fourth species, *P. micropetala*, which grew up to 6 m tall. We did not include this species in our analyses (although its inclusion does not quantitatively affect our results given small sample size), as it is unknown if *H. erato* uses it. Nevertheless, *P. micropetala* is a near relative to *P. biflora* [26] and *Heliconius erato* are known to seasonally switch hosts when their preferred hostplant is depleted [45-48]. Regardless, the presence of these plants in La Barra resonates with the idea that the lowland forest undercanopy offers support for taller *Passiflora* [16, 27]. Further studies could confirm whether *H. e. venus* uses *P. micropetala*, which may relate to its higher flight-height compared to *H. chestertonii*.

Similar to Giraldo-Pamplona, Corrales-Osorio [49], we found that the structure of high and low-altitude forests in Colombia differs such that the former, occupied by *H. chestertonii*, has generally shorter trees at higher densities. This habitat is also colder and drier, with stronger microclimatic fluctuations [22]. These differences mirror those between the habitats of *H. e. cyrbia* and *H. himera* [16] and may result in shared sources of divergent selection across both pairs of taxa. For instance, oviposition behaviour in butterflies is affected by microclimatic conditions [50, 51].

Using dummy models, Merckx, Van Dongen [52] have also showed that *Pararge aegeria* warms up faster when ambient temperatures are higher, experiencing less convective cooling at lower heights (0.9m difference) in forests with a more sheltered structure. In *Drosophila*, pupation-height is also partly driven by desiccation risk, which decreases close to the ground [53, 54]. It is therefore plausible that *H. e. venus* and *H. chestertonii* are flight-height adjusted to compensate for habitat differences influencing microclimate-related traits, rather than to match resource availability.

In conclusion, we showed that two derived mountain specialist *Heliconius* have independently evolved flight behaviours via trait-shifts similar in trajectory, suggesting local adaptations in forests at different elevations. In Colombia, this pattern does not appear to be driven by differences in the vertical distribution of resources. We suggest that these parallel differences may instead relate to effects of microclimatic conditions, variable between forest types. These results add to a limited list of studies exploring potential mechanisms underlying differences in vertical space use between closely related insect taxa.

## Acknowledgements

We are thankful to Oralia Rivas, from Hostal Doña Oralia, for guidance during fieldwork in La Barra. Many thanks also to Denise Dell’Aglio for help incorporating flight-height variation data from Ecuadorian species in our analyses.

## Data accessibility

Supplementary information and raw data can be found in Tables S1-S4 and annotated R scripts. Files available in the Dryad Digital Repository.

## Ethics statement

Fieldwork was conducted under the permit no. 530 issued by the Autoridad Nacional de Licencias Ambientales of Colombia (ANLA).

## Author contributions

SHM and RMM conceptualised the study. DFRS and CHGQ performed the data collection under the supervision and support of CPD and CS. DFRS curated the data. SHM, RMM and DFRS designed the analyses. DFRS performed the analyses and wrote the first draft, which was subsequently edited and approved with contributions of all the authors.

